# Drug-Target Interaction prediction using Multi Graph Regularized Nuclear Norm Minimization

**DOI:** 10.1101/455642

**Authors:** Aanchal Mongia, Angshul Majumdar

## Abstract

The identification of interactions between drugs and target proteins is crucial in pharmaceutical sciences. The experimental validation of interactions in genomic drug discovery is laborious and expensive; hence, there is a need for efficient and accurate in-silico techniques which can predict potential drug-target interactions to narrow down the search space for experimental verification.

In this work, we propose a new framework, namely, Multi Graph Regularized Nuclear Norm Minimization, which predicts the interactions between drugs and proteins from three inputs: known drug-target interaction network, similarities over drugs and those over targets. The proposed method focuses on finding a low-rank interaction matrix that is structured by the proximities of drugs and targets encoded by graphs. Previous works on Drug Target Interaction (DTI) prediction have shown that incorporating drug and target similarities helps in learning the data manifold better by preserving the local geometries of the original data. But, there is no clear consensus on which kind and what combination of similarities would best assist the prediction task. Hence, we propose to use various multiple drug-drug similarities and target-target similarities as multiple graph Laplacian (over drugs/targets) regularization terms to capture the proximities exhaustively.

Extensive cross-validation experiments on four benchmark datasets using standard evaluation metrics (AUPR and AUC) show that the proposed algorithm improves the predictive performance and outperforms recent state-of-the-art computational methods by a large margin.

**Author summary:** This work introduces a computational approach, namely Multi-Graph Regularized Nuclear Norm Minimization (MGRNNM), to predict potential interactions between drugs and targets. The novelty of MGRNNM lies in structuring drug-target interactions by multiple proximities of drugs and targets. There have been previous works which have graph regularized Matrix factorization and Matrix completion algorithms to incorporate the standard chemical structure drug similarity and genomic sequence target protein similarity, respectively. We introduce multiple drug-graph laplacian and target-graph laplacian regularization terms to the standard matrix completion framework to predict the missing values in the interaction matrix. The graph Laplacian terms are constructed from various kinds and combinations of similarities over drugs and targets (computed from the interaction matrix itself). In addition to this, we further improve the prediction accuracy by sparsifying the drug and target similarity matrices, respectively. For performance evaluation, we conducted extensive experiments on four benchmark datasets. The experimental results demonstrated that MGRNNM clearly outperforms recent state-of-the-art methods under three different cross-validation settings, in terms of the area under the ROC curve (AUC) and the area under the precision-recall curve (AUPR).

## Introduction

The field of drug discovery in Pharmaceutical Sciences is plagued with the problem of high attrition rate. The task is to find effective interactions between chemical compounds (drugs) and amino-acid sequences/proteins (targets). This is traditionally done through wet-lab experiments which are known to be costly and laborious. An effective and appropriate alternative to avoid costly failures is to computationally predict the interaction probability. A lot of algorithms have been proposed for DTI (Drug-target interaction) prediction in recent years [1, 2], which use small number of experimentally validated interactions in existing databases such as ChEMBL [3], DrugBank [4], KEGG DRUG [5], STITCH [6] and SuperTarget [7]. Identification of drug-target pairs leads to improvements in different research areas such as drug discovery, drug repositioning, polypharmacology, drug resistance and side-effect prediction [8]. For instance, Drug repositioning [9, 10] (reuse of existing drugs for new indications) may contribute to its polypharmacology (i.e. having multiple therapeutic effects). One of the many successfully repositioned drugs is Gleevec (imatinib mesylate). It was originally thought to interact only with the Bcr-Abl fusion gene associated with leukemia but later, it was found to also interact with PDGF and KIT, eventually leading it to be repositioned to treat gastrointestinal stromal tumors as well [11, 12].

There are three major classes of computational methods for predicting DTI: Ligand-based approaches, Docking based approaches, and Chemogenomic approaches. Ligand-based approaches leverage the similarity between target proteins’ ligands to predict interactions [13]. These approaches use the fact that similar molecules tend to share similar properties and usually bind similar proteins [14]. However, lack of known ligands per protein in some cases might compromise the reliability of results. Docking-based approaches are well-accepted and utilize the 3D structure information of a target protein and a drug; and then run a simulation to estimate the likelihood that they will interact or not [15–17]. But docking is heavily time-consuming and cannot be applied to protein families for which the 3D structure is difficult to predict or is unavailable [18] for example the G-protein coupled receptors (GPCRs).

Chemogenomic approaches overcome the challenges of traditional methods and thus, have recently gained much attention. The approaches under this category work with widely abundant biological data, publicly available in existing online databases and process information (chemical structure graphs and genomic sequences for the drugs and targets) from both the drug and target sides simultaneously for the prediction task. These approaches can further sub-classified based on the representation of the input data: Feature-based methods and Similarity-based methods. Feature-based techniques are machine learning methods, which take their inputs in the form of feature vectors, representing a set of instances (i.e. drug-target pairs) along with their corresponding class labels (i.e. binary values indicating whether or not an interaction exists). Examples of typical feature based methods include Decision Tree (DT), Random Forest (RF) [25] and Support Vector Machines (SVM) to build classification models based on the labeled feature vectors [19]. Positive instances are the known interactions and negative instances, the non-interactions. It should be noted that negative instances here include both non-interactions and unknown drug-target interactions (false negatives). The other category of chemogenomic techniques, Similarity-based methods, use two similarity matrices corresponding to drug and target similarity, respectively, along with an interaction matrix which indicates which pairs of drugs and targets interact.

Let us discuss the similarity between the said DTI problem and the problem of collaborative filtering (CF). CF is a standard problem in information retrieval. It is used in recommendations systems (e.g. in Netflix movie recommendations and Amazon product recommendations). There is a database of user’s and their ratings on items (movies, products, etc.). Obviously, not all the ratings are available; users typically rate only a small subset of items. The objective is to estimate the ratings of all the users on all the items. If that can be done accurately, recommendation accuracy increases. The similarity between DTI and CF should be straightforward now; the drugs play the role of users and the targets play the role of items. The interactions are similar to the ratings. Over the years, many approaches originally developed for CF have been leveraged to solve the DTI problems. In both CF [20] and DTI [21–23], the initial techniques were based on simple neighborhood-based models. In order to predict the interaction of a (active) drug on a target, the first step is to find out similar (neighbor) drugs by computing some kind of a similarity score. Once the neighborhood is obtained, the interaction value from the drugs in the neighborhood are weighted (by the normalized similarity score) to interpolate the interaction of the active drug on the target. The second approach was based on bipartite local models. In such models, a local model is built for every drug and target. For example in [24] an SVM was trained for each to predict the interaction of each drug on all targets and each target on all drugs. Finally, the decision from the two was fused. This is just an example, there are other techniques falling under this generic approach like [25, 26]. The third category is based on network diffusion models. One technique for DTI prediction based on such models is based on a random walk on the network with a predefined transition matrix [27]. Another work falling under this category, predicts interactions by finding simple path (without loops) between nodes of the network. The fourth approach is based on matrix factorization. These techniques were originally developed for collaborative filtering [28]. It is assumed that the drugs and targets are characterized by latent factors. The probability of interaction is high when the latent factors match; i.e. when the inner product has a high value. Therefore, it is logical to express the interaction matrix as a (an inner) product of drug and target latent factors. This allows matrix factorization (and its variants) to be applied [29, 30]. The fifth and final approach is based on classification. The chemical/biological information is used to generate features for drugs and targets individually. The two features are then concatenated and the corresponding interaction is assumed to the class corresponding to this feature. Any standard classifier can be used for the final classification. In such class of techniques, the emphasis is on different feature selection mechanisms [31, 32].

In a very recent review paper [2] it was empirically shown that matrix factorization based techniques yields by far the best results. The fundamental assumption behind matrix factorization to work is that there are very few (latent) factors that are responsible for drug target interactions. This is the reason, one can factor the DTI matrix into a tall (drug) latent factor matrix and a fat (target) latent factor matrix. Mathematically speaking, the assumption is that the DTI matrix is of low-rank. Matrix factorization is being used to model low-rank matrices for the past two decades since the publication of Lee and Seung’s seminal paper [33]. However, matrix factorization is a bi-linear non-convex problem; there are no convergence guarantees. In order to ameliorate this problem, mathematicians proposed an alternate approach based on nuclear norm minimization [34–36]. The nuclear norm is the closest convex surrogate to the rank minimization (known to be NP-hard) problem and there are provable mathematical guarantees on its equivalence to rank minimization.

The standard versions of both the matrix factorization and nuclear norm minimization techniques are unable to incorporate similarity information of the drugs and the targets. In recent studies [30, 37], it was shown that the best results are obtained when these technique incorporate graph regularization penalties into them. But, these works regularize the objective function by taking into account, just the standard chemical structure similarity for drugs (*S*_*d*_) and the genomic sequence similarity for targets (*S*_*t*_). No study in literature gives a clear picture of which kind of similarities would be the best for DTI prediction. We, therefore, incorporate different other kinds of similarities and a combination of them as a multi graph Laplacian regularization with Nuclear Norm Minimization for DTI prediction. The algorithm uses four new similarity measures over the drugs and targets, apart from the standard similarities to construct the graph Laplacians. The four newly incorporated similarities are computed from the interaction matrix and take into account the Cosine similarity, Correlation, Hamming distance and Jaccard similarity between the drugs and targets. To the best of our knowledge, this is the first work on multiple graph laplacian regularized nuclear norm minimization for DTI prediction.

## Materials and methods

### Dataset Description

We use the four benchmark datasets introduced in [21] concerning four different classes of target proteins, namely, enzymes (Es), ion channels (ICs), G protein-coupled receptors (GPCRs) and nuclear receptors (NRs). The data was simulated from public databases KEGG BRITE [38], BRENDA [39] SuperTarget [7] and DrugBank [4].

The data gathered from these databases is formatted as an adjacency matrix, called interaction matrix between drugs and targets, encoding the interaction between n drugs and m targets as 1 if the drug *d*_*i*_ and target *t*_*j*_ are known to interact and 0, otherwise.

Along with the interaction matrix, drug similarity matrix *S*_*d*_ and a target similarity matrix *S*_*t*_ are also provided. In *S*_*d*_, each entry represents the pairwise similarity between the drugs and is measured using SIMCOMP [40]. It represents the chemical structure similarity computed by the number of shared substructures in chemical structures between two drugs. In *S*_*t*_, the similarity score between two proteins is the genomic sequence similarity. It is based on the amino acid sequences of the target protein and is computed using normalized Smith–Waterman [41].

The similarity matrices *S*_*d*_ and *S*_*t*_ constitute the most standard similarities that have been used in the DTI prediction task hitherto. We use these similarities along with the following four more similarities computationally derived from the drug-target interaction matrix to form the graph laplacian terms:

- Cosine similarity: measures the cosine of the angle between two drug/target vectors projected in a multi-dimensional space. Its value ranges from −1 (exactly opposite) to 1 (exactly the same). Given two *n*-dimensional drug/target vectors, the cosine similarity is calculated as follows:

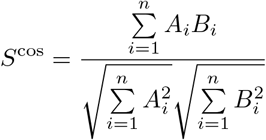
- Correlation: computes the Pearson’s linear correlation coefficient indicating the extent to which two variables are linearly related. It has a value between +1 and −1, where 1 is total positive linear correlation, 0 is no linear correlation, and −1 is total negative linear correlation. For a pair (say A and B) of drugs/targets with sample size *n*, it is given by:

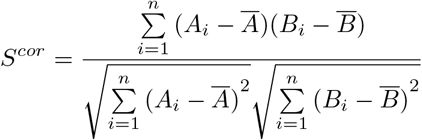

where

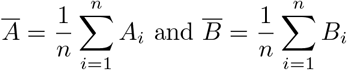
- Hamming similarity: has been computed using Hamming distance. For any two drugs/targets, the hamming distance is the percentage of interaction positions that differ. We calculate Hamming distance based similarity by simply subtracting hamming distance from 1, giving us its complementary (the percentage of common interaction positions for a pair of drugs/targets). It can be calculated as follows:

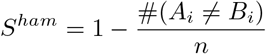
- Jaccard similarity: is defined as the percentage of common non-zero interaction positions for the two given sample sets of drugs/target.

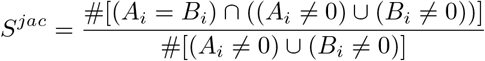

Table 1 summarizes the statistics of all four datasets.

**Table 1.** Drugs, Targets and Interactions in each dataset used for validation.

| Datasets | NR | GPCR | IC | E |
| --- | --- | --- | --- | --- |
| Drugs | 54 | 223 | 210 | 445 |
| Targets | 26 | 95 | 204 | 664 |
| Interactions | 90 | 635 | 1476 | 2926 |

### Nuclear Norm minimization

Let us assume that *X* is the adjacency matrix where each entry denotes interaction between a drug and target (1 if they interact, 0 otherwise). Unfortunately, we only observe this matrix partially because all interactions are not known. If *M* denotes the partially observed adjacency matrix, the mathematical relation between *X* and *M* is expressed as:

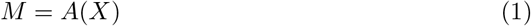

Here *A* is the sub-sampling operator which is element-wise multiplied to *X*. It is a binary mask that has 0’s where the interaction *X* has not been observed or is unknown and 1’s where they have been. Our problem is to recover *X*, given the observations *M*, and the sub-sampling mask *A*. It is known that *X* is of low-rank. Ideally, X should be recovered by (2).

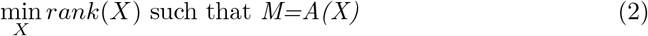

Unfortunately, rank minimization is an NP-hard problem with doubly exponential complexity, therefore solving it directly is not feasible.

Traditionally, a low-rank matrix has been modeled as a product of a thin and a fat matrix and recovered using Matrix Factorization techniques [33]. But, Matrix Factorization is a bi-linear non-convex problem, therefore there is no guarantee for global convergence. In the past decade, mathematicians showed that the rank minimization problem can be relaxed by its convex surrogate (nuclear norm minimization) with provable guarantees [34, 35] This turns out to be a convex problem that can be solved by Semi-Definite Programming. More efficient solvers have also been proposed. Problem (2) is expressed as (3)

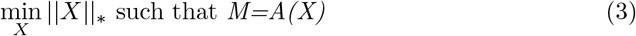

Here the nuclear norm (‖ ‖_*_) is defined as the sum of singular values of data matrix *X*. It is the *l*_1_ norm of the vector of singular values of X and is the tightest convex relaxation of the rank of the matrix, and therefore its ideal replacement.

Here, (3) is a constrained formulation for the noiseless scenario, usually its relaxed version, (4) is solved.

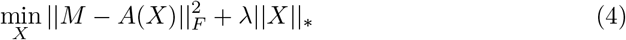

One of the efficient solvers for Nuclear Norm minimization is the Singular value shrinkage (SVS) algorithm [42].

#### Algorithm 1 Singular value shrinkage

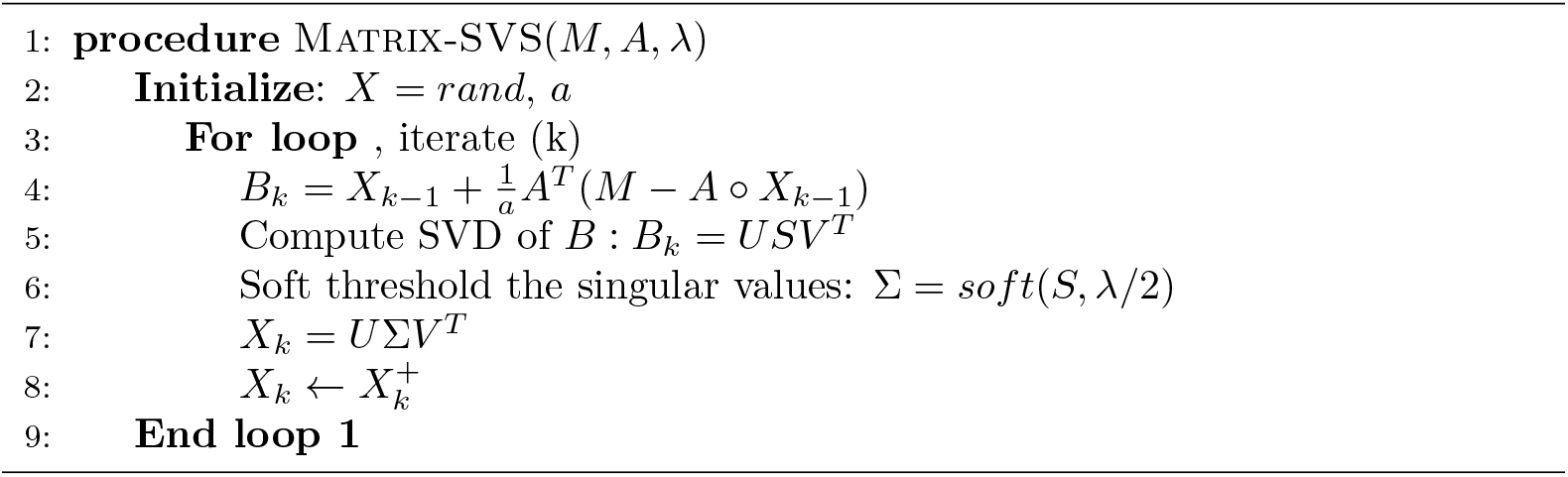

### Multi Graph Regularized Nuclear Norm Minimization

Nuclear Norm based Low-rank Matrix Completion is not our contribution, it has been around since the past decade. The problem with standard Nuclear norm minimization (NNM) is that it cannot accommodate associated information such as Similarity matrices for Drugs and Targets. But, it has been seen in recent studies that accommodating the similarity information is crucial for improving the DTI prediction results. The current works have incorporated the standard similarity measures for drugs and targets in matrix factorization [30] and Matrix completion [37] frameworks. It is imperative that NNM should be capable of taking into account more types and combinations of similarities. To achieve this, we have augmented four other types of similarities between drugs/targets and presented Multi Graph regularized Nuclear Norm Minimization (MGRNNM).

Graph regularization assumes that points close to each other in the original space should also be close to each other in the learned manifold (**Local Invariance assumption**). So, Graph regularization would allow the algorithm to learn manifolds for the drug and target spaces in which the data is assumed to lie.The multi graph regularized version of Nuclear norm minimization, aims to prevent over fitting and greatly enhance the generalizing capabilities. It is incorporated into the formulation/objective function as Laplacian weights corresponding to drugs and targets:

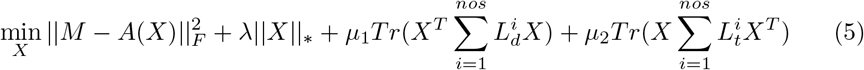

where *λ* ≥ 0, *μ*_1_ ≥ 0 and *μ*_2_ ≥ 0 are parameters balancing the reconstruction error of NNM in the first two terms and graph regularization in the last two terms, *Tr*(.) is the trace of a matrix, *nos* stands for number of similarity matrices (*nos* = 5 in our case).

If, say we consider a single similarity matrix for drugs (*S*_*d*_) and that for targets (*S*_*t*_), then *L*_*d*_ = *D*_*d*_ − *S*_*d*_ and *L*_*t*_ = *D*_*t*_ − *S*_*t*_ are the graph Laplacians [43] for *S*_*d*_ (drug similarity matrix) and *S*_*t*_ (target similarity matrix), respectively, and 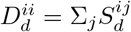 and 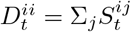 are degree matrices.

Problem (5) is solved using a variable splitting approach [44]. The augmented Lagrangian is expressed as (6). We introduce two new proxy variables *Z* and *Y* such that *Z*^*T*^ = *X* and *Y* = *X*.

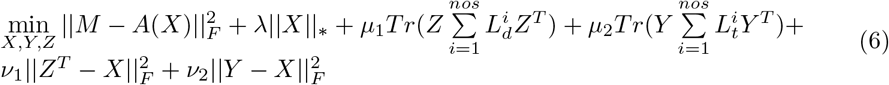

The variables are updated using ADMM [45, 46]. This leads to the following subproblems (7), (8) and (9)

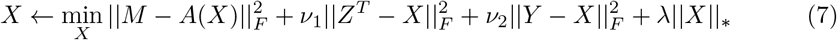

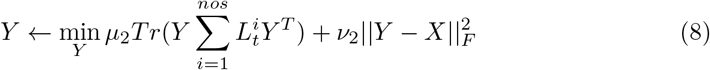

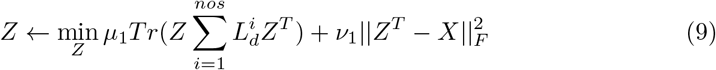

Problem (7) can be expressed as a standard NNM probelm (by column stacking the variables).

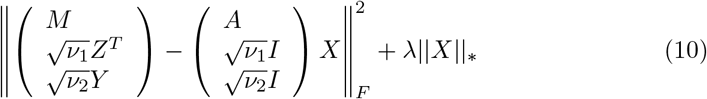

To solve for *Y* and *Z*, we differentiate (8) and (9) wrt *Y* and *Z*, respectiveley.

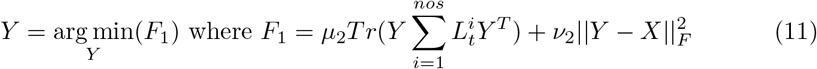

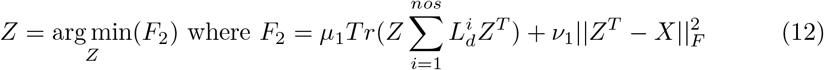

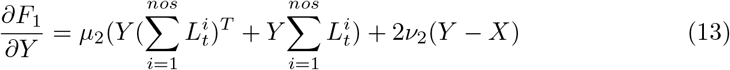

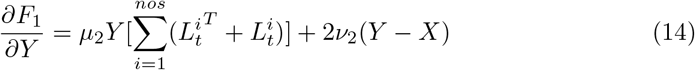

Since *L*_*t*_ is a symmetric matrix, 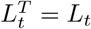. So,

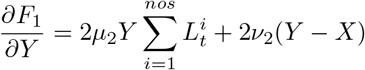

Equating the derivative to zero, we get:

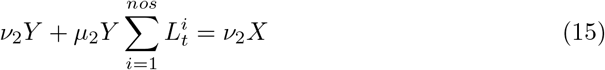

The matrix equation of this form (AT+TB=C) cannot be solved directly for variable T and is called Sylvester equation. Such an equation has a unique solution when the eigenvalues of A and -B are distinct.

A similar Sylvester equation and update step for *Z* can be obtained by differentiating *F*_2_ and equating to 0.

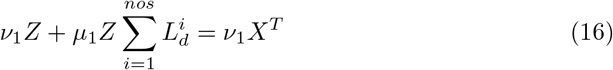

It can be shown that computing the sum of the Graph Laplacians is equivalent to computing the Laplacian from the sum of various similarity matrices involved. For instance, consider the sum of drug Graph Laplacians:

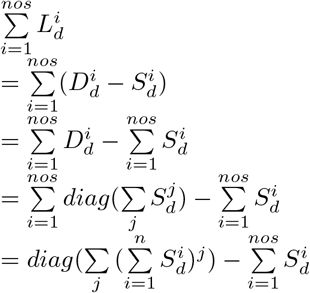

Let 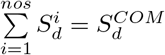 where 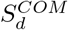 stands for combined similarity for drugs. Essentially,

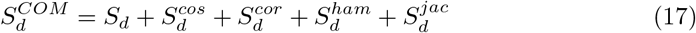

łThen,

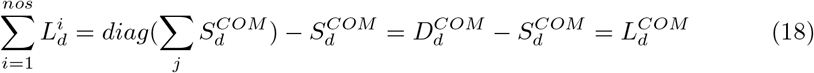

Here, 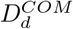 and 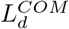 denote combined degree matrix and combined Laplacian matrix (sum of graph laplacians) for drugs. Of note, the individual Laplacians or the similarities can be weighted unequally to give more or less emphasis on a specific type of similarity. The pseudo-code for MGRNNM has been given in Algorithm 2.

#### Algorithm 2 Multi Graph regularized Nuclear Norm Minimization

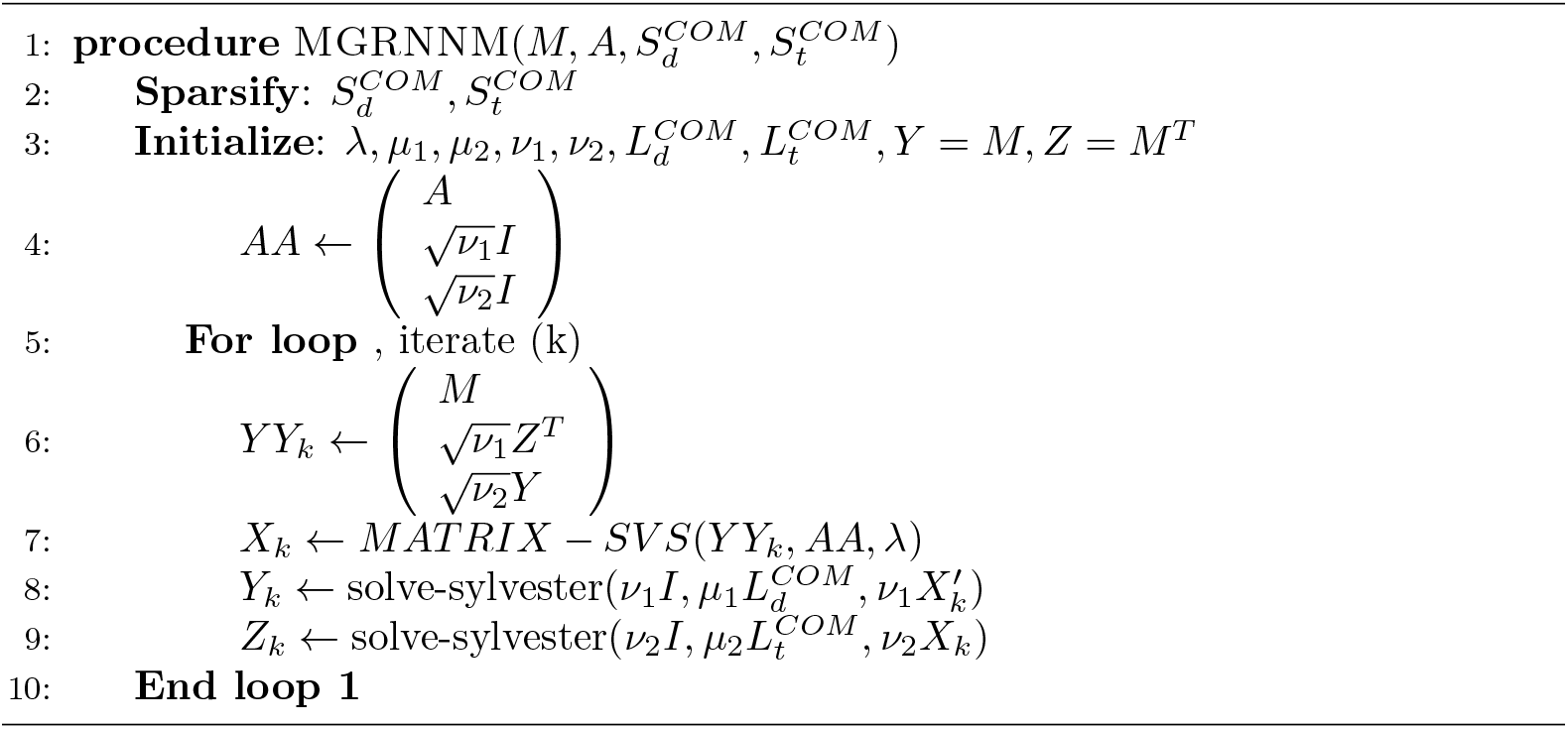

The standard NNM is a convex problem and the introduced graph regularization penalties are also convex, so entire formulation (5), being a sum of convex functions, is convex. Therefore it is bound to converge to a global minima. We chose the number of iterations such that the algorithm halts when the objective function does not change with iterations. A sample convergence plot for one of the datasets for drug-target pair prediction has been shown in Figure 1

**Fig 1.**
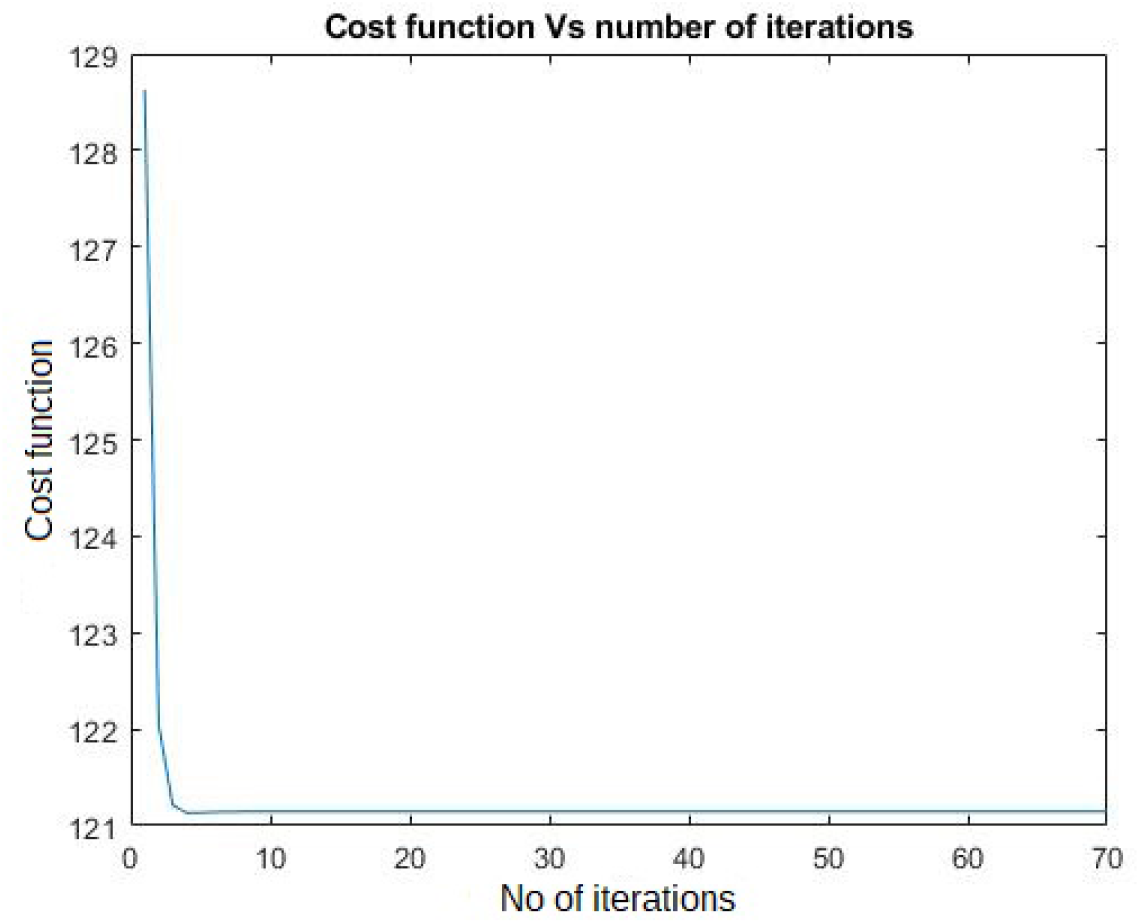
Converge plot of the MGRNNM algorithm for NR dataset with cross validation setting CVS1 (drug-target pair prediction).

### Time Complexity of MGRNNM

The algorithm is iterative, so we only discuss the time complexity per iteration. In each iteration, we solve the two Sylvester equations and one NNM. NNM itself is an iterative algorithm that requires solving Singular value decomposition in each iteration, the order of complexity of which is *O*(*n*^3^). The complexity of solving Sylvester equation is *O*(*n*^*b*^*.log*(*n*)) [47] where *b* is between 2 and 3.

## Results and Discussion

### Experimental Setup

We validated our proposed method by comparing it with recent and well-performing prediction methods proposed in the literature. Out of the 5 approaches with which we compare,

- Three are specifically designed for DTI task (WGRMF: Weighted Graph Regularized Matrix Factorization, CMF: Collaborative Matrix Factorization, RLS WNN: Regularized Least square Nearest neighbor profile) [30, 48, 49];
- One being traditional matrix completion (MC: matrix completion) [42] and
- Last one being a naive solution to our problem, available as an unpublished work (MCG: matrix completion on graphs). Of note, the Space complexity of MCG is *O*(*n*^4^) while that of MGRNNM is *O*(*n*^2^). [50])

All baselines designed for DTI problem are recent and are already compared against older methods.

We performed 5 repetitions of 10-fold cross-validation (CV) for each of the methods under three cross-validation setting (CVS) [2]:

- CVS1/Pair prediction: random drug–target pairs are left out as the test set for prediction. It is the conventional setting for validation and evaluation.
- CVS2/Drug prediction: entire drug profiles are left out to be used as test set. It tests the algorithm’s ability to predict interactions for novel drugs i.e. drugs for which no interaction information is available.
- CVS3/Target prediction: entire target profiles are left out to be used as test set. It tests the algorithm’s ability to predict interactions for novel targets.

In 10-fold CV, data was divided into 10 folds and out of those 10 folds, one was left out as the test set while the remaining 9 folds were treated as the training set. As the evaluation metrics, We used:

- AUC: AUC stands for “Area under the ROC Curve.” That is, AUC measures the entire two-dimensional area underneath the entire ROC curve (a plot showing the true positive rate for a method as a function of the false positive rate). AUC provides an aggregate measure of performance across all possible classification thresholds. One way of interpreting AUC is the probability that the model ranks a random positive example more highly than a random negative example. The higher it is, the better the model is.
- AUPR: We also evaluated the performance by AUPR (Area Under the Precision-Recall curve), because AUPR punishes highly ranked false positives much more than AUC, this point being important practically since only highly ranked drug-target pairs in prediction will be biologically or chemically tested later in an usual drug discovery process, meaning that highly ranked false positives should be avoided. The precision-recall curve shows the tradeoff between precision and recall for different thresholds. A high area under the curve represents both high recall and high precision, where high precision relates to a low false positive rate, and high recall relates to a low false negative rate. High scores for both show that the classifier is returning accurate results (high precision), as well as returning a majority of all positive results (high recall).

### Preprocessing

Each of the drug and target similarity matrices were summed up to compute the combined similarity matrices 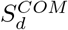 and 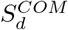 (equation (17)). The combined similarity matrices were further sparsified by using p-nearest neighbor graph which is obtained by keeping only the similarity values of the nearest neighbors for each drug/target in the similarity matrices. The usage of such a pre-processing, as shown by [30], helps learn a manifold on or near to which the data is assumed to lie which, in turn, is expected to preserve the local geometries of the original data and hence give more accurate results.

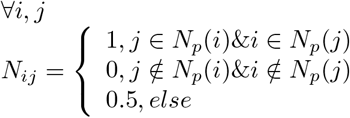

where *Np*(*i*) is the set of *p* nearest neighbors to drug *d*_*i*_. Similarity matrix sparsification is done by element-wise multiplying it with *N*_*ij*_. In the next step, the combined graph laplacian terms are computed. Also, instead of graph laplacians (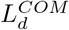 and 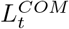), we have used normalized graph laplacians (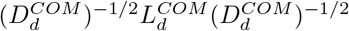 and 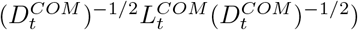 instead as normalized graph Laplacians are known to perform better in many cases [51].

### Parameter settings

For setting the parameters of our algorithm, we performed cross-validation on the training set on the parameters *p, λ, μ*_1_, *μ*_2_, *ν*_1_, *ν*_2_ to find the best parameter combination for each dataset, under each cross-validation setting. As mentioned earlier, the individual laplacians or the similarities can be weighted unequally to give more or less emphasis on a specific type of similarity, we weigh the Cosine, Correlation and Jaccard similarities heavily (4 times) relative to Hamming similarity. This was done because hamming similarity showed the least improvement in prediction accuracy as compared to the other three similarities when taken into account along with standard similarities (Refer Figure 2). For the other methods, we to set the parameters to their optimal (which were found to be already optimal) in [2].

**Fig 2.**
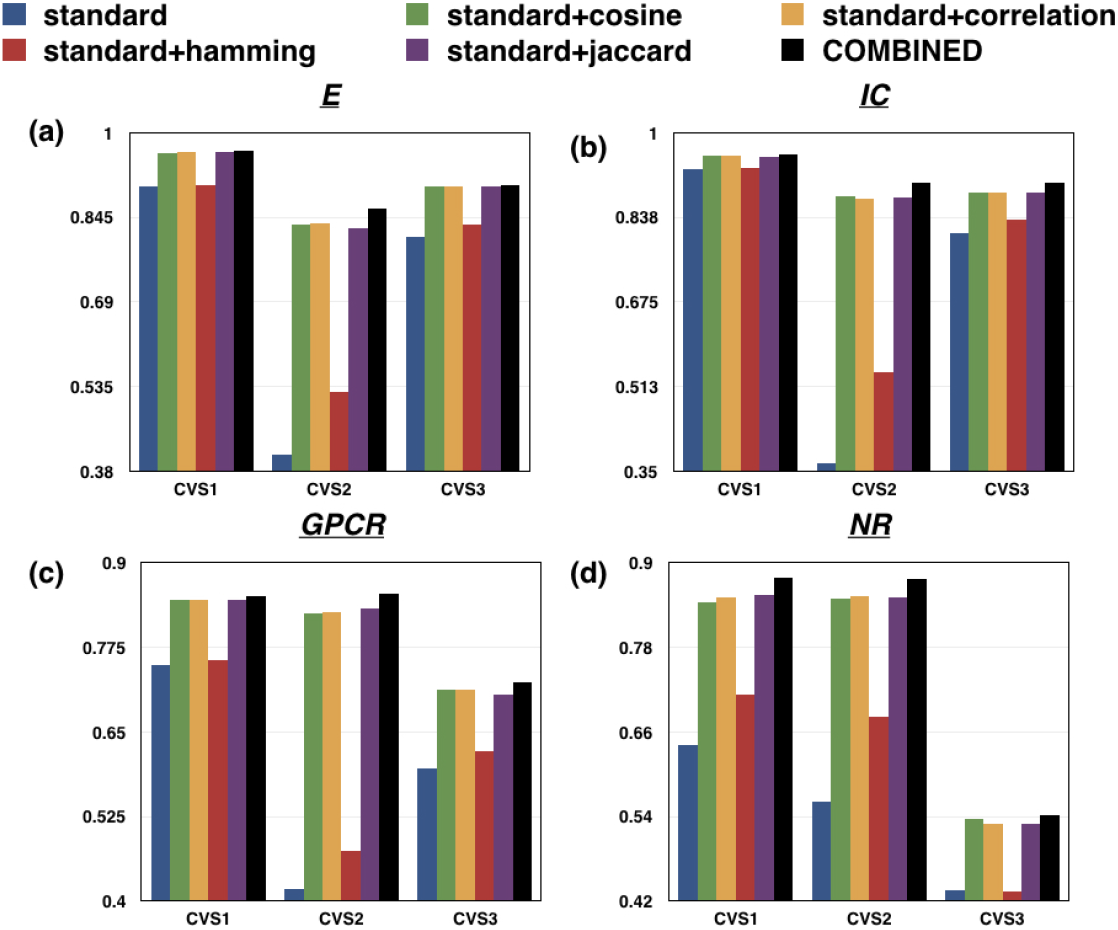
Bar plots depicting that incorporating all the similarities for drugs and targets for prediction task yields best results for every dataset **(a)** E **(b)** IC **(c)** GPCR and **(d)** NR under the three cross-validation settings in comparison to the cases where each type of similarity was considered separately. Here, “standard” represents the case when only the chemical structure similarity for drugs and genomic sequence similarity for targets were taken into account and “COMBINED” refers to the use case where all the similarity matrices (standard similarity, Cosine similarity, Correlation, Hamming similarity and Jaccard similarity) were considered.

### Interaction Prediction

Tables 2, 4 and 6 show the AUPR results and tables 3, 5 and 7 show the AUC results from the above-mentioned cross validation settings. MGRNNM outperforms the state-of-the-art prediction methods. The second column in each table shows the results of our algorithm when only the standard similarity matrices (*S*_*d*_: chemical structure similarity for drugs, *S*_*t*_: Genomic sequence similarity for target proteins) were used for prediction.

**Table 2.**
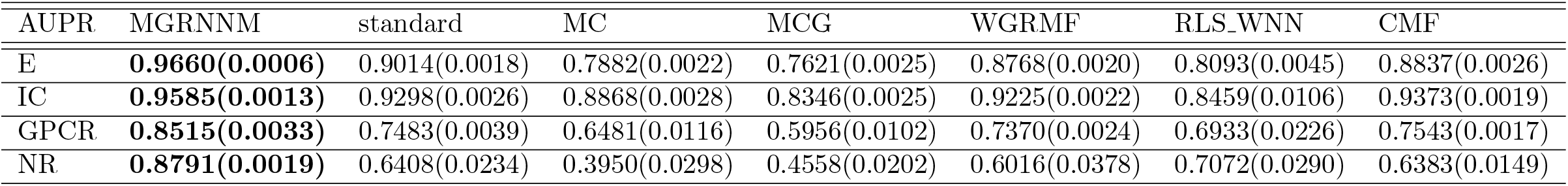
AUPR results for interaction prediction under validation setting CVS1.

| AUPR | MGRNNM | standard | MC | MCG | WGRMF | RLS_WNN | CMF |
| --- | --- | --- | --- | --- | --- | --- | --- |
| E | **0.9660(0.0006)** | 0.9014(0.0018) | 0.7882(0.0022) | 0.7621(0.0025) | 0.8768(0.0020) | 0.8093(0.0045) | 0.8837(0.0026) |
| IC | **0.9585(0.0013)** | 0.9298(0.0026) | 0.8868(0.0028) | 0.8346(0.0025) | 0.9225(0.0022) | 0.8459(0.0106) | 0.9373(0.0019) |
| GPCR | **0.8515(0.0033)** | 0.7483(0.0039) | 0.6481(0.0116) | 0.5956(0.0102) | 0.7370(0.0024) | 0.6933(0.0226) | 0.7543(0.0017) |
| NR | **0.8791(0.0019)** | 0.6408(0.0234) | 0.3950(0.0298) | 0.4558(0.0202) | 0.6016(0.0378) | 0.7072(0.0290) | 0.6383(0.0149) |

**Table 3.** AUC results for interaction prediction under validation setting CVS1.

| AUPR | MGRNNM | standard | MC | MCG | WGRMF | RLS_WNN | CMF |
| --- | --- | --- | --- | --- | --- | --- | --- |
| E | **0.9955(0.0003)** | 0.9798(0.0004) | 0.8753(0.0023) | 0.9596(0.0015) | 0.9647(0.0013) | 0.9635(0.0014) | 0.9705(0.0013) |
| IC | **0.9947(0.0004)** | 0.9829(0.0012) | 0.9415(0.0015) | 0.9539(0.0010) | 0.9747(0.0022) | 0.9786(0.0026) | 0.9832(0.0008) |
| GPCR | **0.9785(0.0020)** | 0.9531(0.0028) | 0.8110(0.0055) | 0.8977(0.0047) | 0.9432(0.0010) | 0.9458(0.0044) | 0.9493(0.0031) |
| NR | **0.9660(0.0056)** | 0.9083(0.0058) | 0.5882(0.0253) | 0.8315(0.0165) | 0.8892(0.0153) | 0.9329(0.0114) | 0.8679(0.0124) |

**Table 4.**
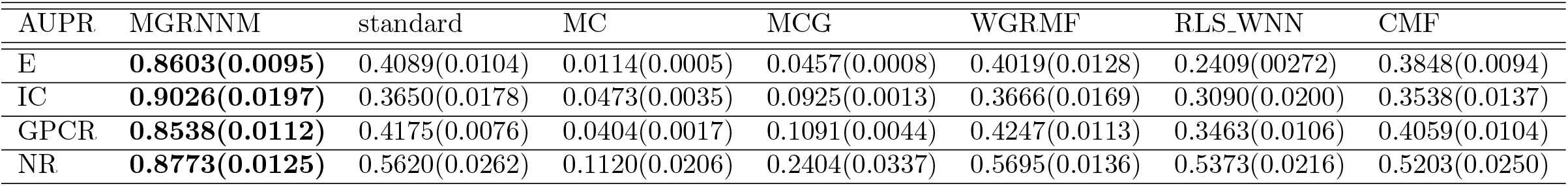
AUPR results for interaction prediction under validation setting CVS2.

**Table 5.** AUC results for interaction prediction under validation setting CVS2.

| AUPR | MGRNNM | standard | MC | MCG | WGRMF | RLS_WNN | CMF |
| --- | --- | --- | --- | --- | --- | --- | --- |
| E | **0.9460(0.0033)** | 0.8260(0.0108) | 0.5060(0.0090) | 0.7413(0.0118) | 0.7982(0.0144) | 0.7755(0.0093) | 0.7952(0.0110) |
| IC | **0.9714(0.0095)** | 0.7913(0.0090) | 0.5512(0.0034) | 0.7196(0.0071) | 0.7902(0.0149) | 0.7669(0.0140) | 0.7576(0.0125) |
| GPCR | **0.9567(0.0084)** | 0.8805(0.0024) | 0.5855(0.0039) | 0.7745(0.0027) | 0.8800(0.0025) | 0.8524(0.0072) | 0.8067(0.0067) |
| NR | **0.9533(0.0127)** | 0.8452(0.0215) | 0.5294(0.0200) | 0.6992(0.0244) | 0.8615(0.0244) | 0.8390(0.0261) | 0.8124(0.0228) |

**Table 6.** AUPR results for interaction prediction under validation setting CVS3.

| AUPR | MGRNNM | standard | MC | MCG | WGRMF | RLS_WNN | CMF |
| --- | --- | --- | --- | --- | --- | --- | --- |
| E | **0.9041(0.0125)** | 0.8087(0.0156) | 0.0124(0.0005) | 0.0691(0.0009) | 0.8070(0.0185) | 0.5465(0.0144) | 0.7808(0.0131) |
| IC | **0.9029(0.0024)** | 0.8079(0.0096) | 0.0421(0.0043) | 0.2256(0.0038) | 0.8128(0.0069) | 0.7437(0.0088) | 0.7786(0.0108) |
| GPCR | **0.7228(0.0323)** | 0.5963(0.0336) | 0.0549(0.0105) | 0.1061(0.0027) | 0.6093(0.0314) | 0.5397(0.0193) | 0.5989(0.0323) |
| NR | **0.5418(0.0309)** | 0.4356(0.0177) | 0.0850(0.0227) | 0.2669(0.0288) | 0.4643(0.0183) | 0.4907(0.0326) | 0.4774(0.0173) |

**Table 7.** AUC results for interaction prediction under validation setting CVS3.

| AUPR | MGRNNM | standard | MC | MCG | WGRMF | RLS_WNN | CMF |
| --- | --- | --- | --- | --- | --- | --- | --- |
| E | **0.9683(0.0043)** | 0.9246(0.0091) | 0.5234(0.0057) | 0.8065(0.0012) | 0.9338(0.0071) | 0.9067(0.0105) | 0.9272(0.0050) |
| IC | **0.9541(0.0019)** | 0.9346(0.0041) | 0.4724(0.0065) | 0.7871(0.0069) | 0.9460(0.0034) | 0.9286(0.0046) | 0.9368(0.0032) |
| GPCR | **0.8975(0.0093)** | 0.8798(0.0134) | 0.5683(0.0310) | 0.6289(0.0151) | 0.8892(0.0110) | 0.8694(0.0146) | 0.8966(0.0073) |
| NR | 0.7502(0.0285) | 0.7263(0.0211) | 0.3767(0.0204) | 0.6522(0.0297) | 0.7967(0.0132) | 0.8124(0.0202) | **0.8373(0.0083)** |

### Validation of Multiple similarities

To precisely analyze the consequence of multiple similarities incorporation, we observed the mean AUPR for several cases:

- standard: When only the standard similarity matrices (*S*_*d*_: chemical structure similarity for drugs, *S*_*t*_: Genomic sequence similarity for target proteins) were used for prediction.
- standard+Cosine: When Cosine similarity between each pair of drugs/targets (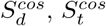) was taken into account along with standard similarities.
- standard+Correlation: When Pearson’s linear Correlation between each pair of drugs/targets (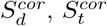) was taken into account along with standard similarities.
- standard+Hamming: When Hamming similarity between each pair of drugs/targets (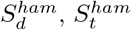) was taken into account along with standard similarities.
- standard+Jaccard: When Jaccard similarity between each pair of drugs/targets (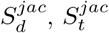) was taken into account along with standard similarities.
- COMBINED: When all five similarity types between each pair of drugs/targets (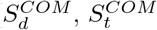) were taken into account.

The analysis was carried out for every dataset under all the three cross-validation settings. Figure 2 clearly depicts that incorporating all the similarities for drugs and targets for prediction task yields the best results.

## Conclusion

Drug-target interaction prediction is a crucial task in genomic drug discovery. Many computational techniques have been proposed in the literature. In this work, we presented a novel chemogenomic approach for predicting the drug-target interactions, MGRNNM (Multi Graph regularized Nuclear Norm Minimization). It is a graph regularized version of the traditional Nuclear Norm Minimization algorithm which incorporates multiple Graph Laplacians over the drugs and targets into the framework for an improved interaction prediction. The algorithm is generic and can be used for prediction in protein-protein interaction [52], RNA-RNA interaction [53], etc.

The evaluation was performed using three different cross-validation settings, namely CVS1 (random drug-target pairs left out), CVS2 (entire drug profile left out) and CVS3 (entire target profile left out) to compare our method with 5 other state-of-the-art methods (three specifically designed for DTI prediction). In almost all of the test cases, our algorithm shows the best performance, outperforming the baselines. This work can be extended by accounting for more types of drug and target similarities which could be either chemically/biologically driven or obtained from the metadata itself to improve the prediction accuracy even further.

